# Intracellular Optical Doppler Phenotypes of Chemosensitivity in Human Epithelial Ovarian Cancer

**DOI:** 10.1101/2020.09.14.296863

**Authors:** Zhe Li, Ran An, Wendy M. Swetzig, Margaux Kanis, Nkechiyere Nwani, John Turek, Daniela Matei, David Nolte

## Abstract

Development of an assay to predict response to chemotherapy has remained an elusive goal in cancer research. We report a phenotypic chemosensitivity assay for epithelial ovarian cancer based on Doppler spectroscopy of infrared light scattered from intracellular motions in living three-dimensional tumor biopsy tissue measured *in vitro*. The study analyzed biospecimens from 20 human patients with epithelial ovarian cancer. Matched primary and metastatic tumor tissues were collected for 3 patients, and an additional 3 patients provided only metastatic tissues. Doppler fluctuation spectra were obtained using full-field optical coherence tomography through off-axis digital holography. Frequencies in the range from 10 mHz to 10 Hz are sensitive to changes in intracellular dynamics caused by platinum-based chemotherapy. Metastatic tumor tissues were found to display a biodynamic phenotype that was similar to primary tissue from patients who had poor clinical outcomes. The biodynamic phenotypic profile correctly classified 90% [88% to 91% c.i.] of the patients when the metastatic samples were characterized as having a chemoresistant phenotype. This work suggests that Doppler profiling of tissue response to chemotherapy has the potential to predict patient clinical outcomes based on primary, but not metastatic, tumor tissue.

## Introduction

The tumor microenvironment plays an essential role in the complex biological and molecular communication between cancer cells and the host, determining both tumor progression and response to therapy. The microenvironmental influences on the cancer state are associated with mechano-transduction^1, 2^, paracrine signaling, as well as immune cell infiltration and endocrine signaling. Chemosensitivity assays^3, 4^ seek to measure the sensitivity of patient-derived cells to a range of chemotherapies. However, the conventional assays destroy the microenvironmental influences by disaggregating cells from tumor biopsies and growing them in two-dimensional cell culture or as xenografts implanted in host animals. The growth in the environment of the cell culture plate or the animal host changes the cellular phenotype, which may no longer represent the phenotype of the intact tumor. In consequence, chemosensitivity assays have limited ability to test cancer cells from clinical specimens, they lack predictive power for subsequent clinical applications^5-7^, and they rely exclusively on epithelial tumor components. Therefore, a need exists for a phenotypic assay that maintains the three-dimensional microenvironment and tests the functional response of living tissue to selected therapies.

To meet this need, a Doppler fluctuation spectroscopy approach to chemosensitivity testing, called biodynamic imaging (BDI), was developed as the first coherence-domain imaging technique to use intracellular motion as functional image contrast^8^. Intracellular motion includes molecular-motor-dependent transport of vesicles and mitochondria, intranuclear alterations associated with pre- and post-mitotic processes, cytoplasmic streaming, cytoskeletal restructuring, active membrane modulations, and cell shape changes ^9^. These motions range in speed from nanometers per second for cell-scale motions to microns per second for organelle and vesicle transport, generating Doppler frequency shifts from 10 mHz to 10 Hz, respectively. There has been growing recognition of the importance of intracellular dynamics for functional imaging on intact tissue ^10-16^. Biodynamic imaging is a form of full-frame optical coherence tomography (FF-OCT) ^17, 18^ based on off-axis digital holography that uses principles of coherent laser ranging that can quantify the dynamic response of tumors to chemotherapy treatment^19^. Cellular motions are unusual but specific biomarkers of cellular health and response. By penetrating volumetrically into tissue up to 1 mm deep, BDI maps out heterogeneous tissue layers. BDI has previously been applied to drug screening ^20-22^, phenotypic profiling ^23^, and preclinical chemosensitivity testing on ovarian xenografts in mice and canine B-cell lymphoma^19, 24, 25^. Light scattering of near-infrared light from living *ex vivo* tissue biopsies displays Doppler frequency shifts caused by intracellular motion^9^. Chemotherapy agents applied to the living biopsies *in vitro* modify the intracellular dynamics and the associated Doppler frequencies. The Doppler frequency shifts and their changes are interpretable through the speeds of intracellular motions affected by anti-cancer drugs. In a previous study^24^, we used biodynamic imaging to assess ovarian xenografts grown in mice from human ovarian cancer cell lines whose platinum resistance and sensitivity were associated with biodynamic signatures. The work presented here is the first application of BDI to naturally-occurring cancer in human patients.

## Results

### Biodynamic Spectra of Human Ovarian Cancer Biopsies

The core optical components of the biodynamic imaging system are illustrated in Fig. 1a. The system is configured as a Mach-Zehnder interferometer with low-coherence digital holography. Fluctuation spectra of a living tissue sample are obtained from time series of dynamic speckle images on the digital camera, and changes in the spectra are tracked after a drug treatment is applied to the sample well (described in the Methods section). The time-frequency format of a typical drug-response spectrogram is shown in Fig. 1b. Frequency is along the horizontal axis spanning from 10 mHz to 10 Hz. Time is along the vertical axis spanning 17.8 hours: 5.5 hours of baseline followed by 12.3 hours after a treatment is dispensed into the well. The baseline is used for reference, and the treatment is applied at the time of the horizontal blue line. The shifts in the spectral content caused by the drug action are captured in color in the figure, blue representing loss of spectral density and red representing an increase of spectral density. The Doppler frequency is related to the intracellular speed through Δω= qv, where q = 4πn/λ_0_, λ_0_ is the free-space wavelength, n is the refractive index, and v is the internal speed. For reference, a speed of 1 micron per second produces a Doppler frequency shift of 3 Hz at a wavelength of 840 nm in a backscattering geometry. The high-frequency region corresponds to the organelle transport band ^26^. The low-frequency region represents slow membrane rearrangement and possible cell motility ^27, 28^ (described in Table 2 in Methods).

**Figure 1.**
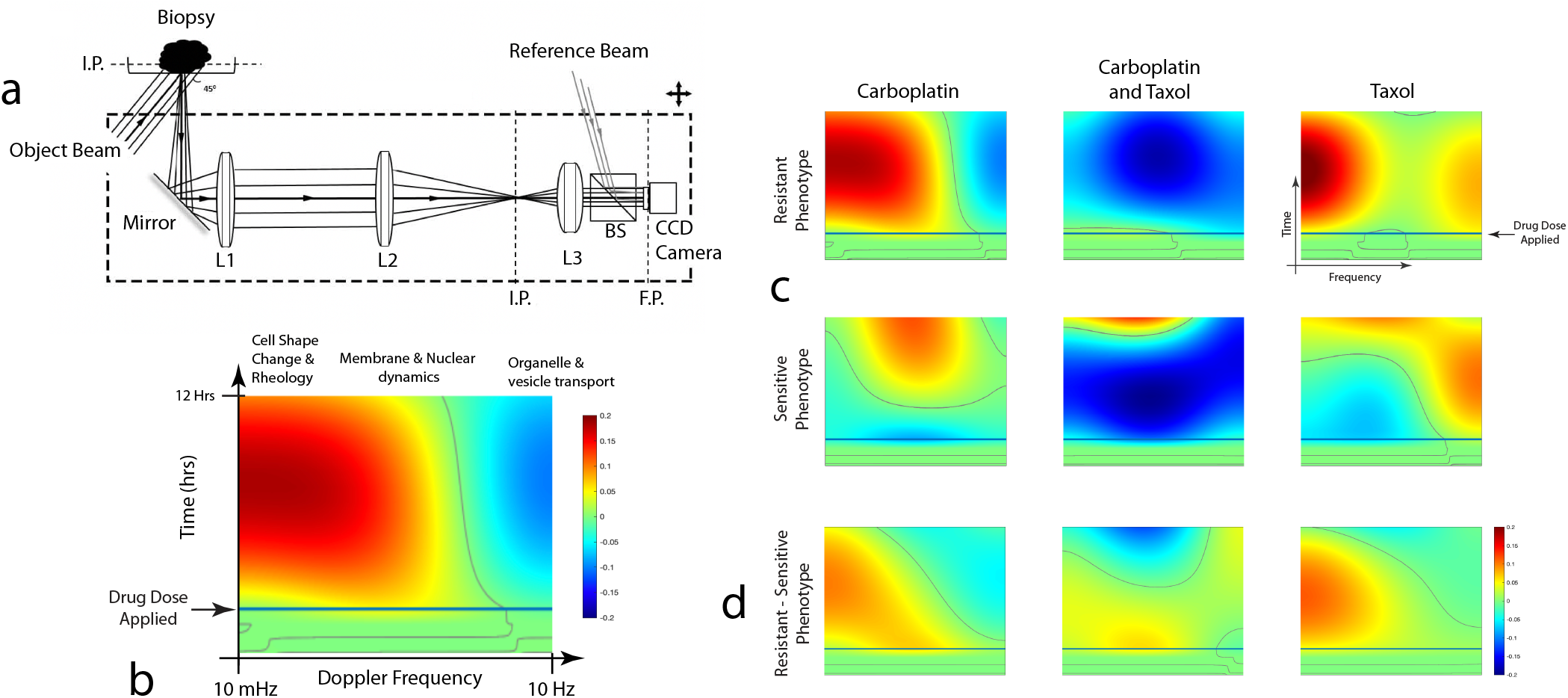
Drug-response spectrograms for poly-lysine-immobilized biopsies treated with carboplatin (25 µM), paclitaxel (5 µM) and carboplatin + paclitaxel (25 µM+5 µM). The axes are the same for all spectrograms. a) A schematic of the biodynamic platform (BDP). The imaging system (including the light source, lenses, beam splitters and the CCD) is placed on an optical platform mounted on a motorized stage that moves in the horizontal plane. (IP image plane. L1-3 lenses. BS beam splitter. FP Fourier plane. CCD charge-coupled device digital camera.) b) In the spectrogram time-frequency format the Doppler frequency spans three orders of magnitude. The spectrogram is the relative change of spectral density relative to the pre-dose baseline. The spectral response is monitored for 12 hours after the dose. c) The average spectrograms (DMSO-subtracted) for resistant and sensitive phenotypes. d) The difference of the resistant spectrograms minus the sensitive.

To generate spectral fingerprints of sensitive versus resistant patients, we partitioned the patient samples into sensitive and resistant groups. Platinum-sensitive tumors were defined as those tumors that did not recur for more than 6 months, while platinum-resistant tumors were those that progressed in less than 6 months after completion of platinum-based therapy. Average drug-response spectrograms in Fig. 1c were obtained for the two cohorts averaged over samples immobilized by poly-lysine (see Methods and Supplemental Information for sample immobilization methods). The differences of the resistant minus the sensitive spectrograms are shown in Fig. 1d with enhanced low frequencies (red shifts) in the resistant samples relative to the sensitive. A so-called “red shift” (increased spectral density at low frequencies and decreased spectral density at high frequency) represents a decrease in average cellular speeds.

### Association of Biodynamic Phenotype with Patient Outcome

The time-frequency representation of the drug-response spectrograms was converted using linear filters into quantitative biomarkers that have strong co-dependence. The features are called “biomarkers” by analogy with genetic or proteomic biomarkers because they represent distinct behavior of the tissue, but the biodynamic biomarkers have not yet been related to specific traditional biomarkers or pathways. Principal component analysis (PCA) was used to find a set of orthogonal biomarkers that are linear combinations of the original biomarkers. The principal components of the spectrogram-based biomarkers having the largest signal-to-noise ratios that differentiate the resistant/sensitive groups are shown in Fig. 2a). The biomarkers are designated by the principal component and by the treatment. For instance, BM2tax+carb represents the 2nd principal component in the singular-value decomposition of the drug response under combination treatment. The feature vectors shown in Fig. 2a) are BM7carb, BM2tax+carb, BM4tax+carb, BM7tax+carb and BM1tax. The feature definitions, selection, and the singular value decomposition (SVD) coefficients of the linear combination of these biomarkers in terms of the original biomarkers are given in Methods.

**Figure 2.**
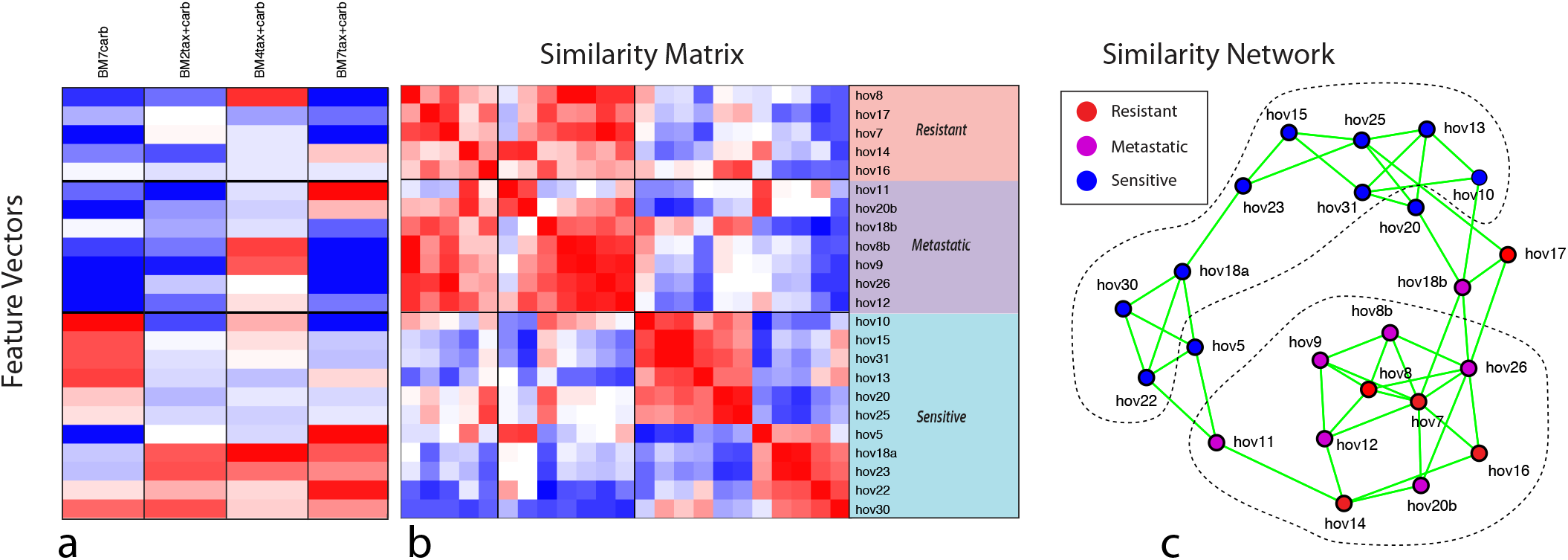
Associating biodynamic features with patient clinical outcomes. a) Feature vectors selected using SVD that show the strongest correlation with clinical outcomes. The patients are partitioned into a resistant group of patients, a metastatic group and a sensitive group. b) The similarity matrix generated from the feature vectors. The matrix is approximately block diagonal. c) The similarity network constructed from the similarity matrix. The sensitive group (red) tends to split into two sub-phenotypes. The metastatic samples share strong similarity with the resistant phenotype, even if the patient was sensitive to treatment.

The selected feature vectors of Fig. 2a) are the central data structure for all downstream machine learning algorithms. The goal is to identify which patients share similarities with each other, and with the sensitive/resistant phenotypes. For instance, the feature vectors are used to construct the similarity matrix in Fig. 2b). The order of the patients was preselected according to their clinical outcomes, separated into resistant, metastatic and sensitive groups. Identical vectors have vector contrast near unity (red), opposite vectors have vector contrast near negative unity (blue), and independent vectors have vector contrast near zero (white). The similarity matrix has an approximately block-diagonal structure. A key observation is that the resistant and metastatic block of biopsies share strong similarities with each other, implying that the metastatic tissues have a dynamic phenotype that is similar to the resistant primary tumor tissue. Because none of the metastatic data were used in the feature selection or training, this resistant phenotype of the metastatic samples is one of the principal conclusions of this study. Among the samples from patients who had sensitive clinical outcomes there appear to be two sub-blocks.

Network theory provides analysis techniques for identifying relationships among a set of feature vectors. A similarity network for this clinical study is shown in **Fig. 2c**. Links in the network are assigned according to a k-neighbor adjacency matrix with k = 3. The patients are color-coded according to their clinical outcomes. Dark red nodes are primary tumors that have resistant clinical outcomes, dark blue nodes are primary tumors that have sensitive clinical outcomes, purple are metastatic tissues. The metastatic samples cluster closely with the resistant primary samples. The network structure provides direct visualization of the similarity matrix. The upper blue group in the network is from the lower right group of sensitive patients in the similarity matrix. The similarity matrix and the network structure suggests two general phenotypes: R-class that includes the resistant and metastatic samples (defined by tumor location and patient clinical outcome), and S-class that contains the sensitive non-metastatic samples.

We used several binary classifier algorithms that were combined to yield an average ensemble chemosensitivity prediction for each patient. These algorithms are: 1) a single-neuron perceptron (logistic regression), 2) a continuous-valued recurrent neural network, 3) log-likelihood and 4) a binary network analysis. These are combined into an ensemble average to predict the chemosensitivity of a patient by using one-hold-out cross-validation for the non-metastatic poly-lysine-immobilized samples. The trained algorithm was then used to predict independently the test set of agar-immobilized samples as well as the metastatic samples. The ensemble average of the classifiers is shown in **Fig. 3a**. The patient sequence in the figure was pre-ordered into resistant, metastatic and sensitive patients and was clustered by similarity within each of those groups. The error bars are the standard errors obtained from alternative training subsets (described in the Supplemental Material (see **Fig. S4**)) that included agar-immobilized training samples as well as the poly-lysine.

**Figure 3.**
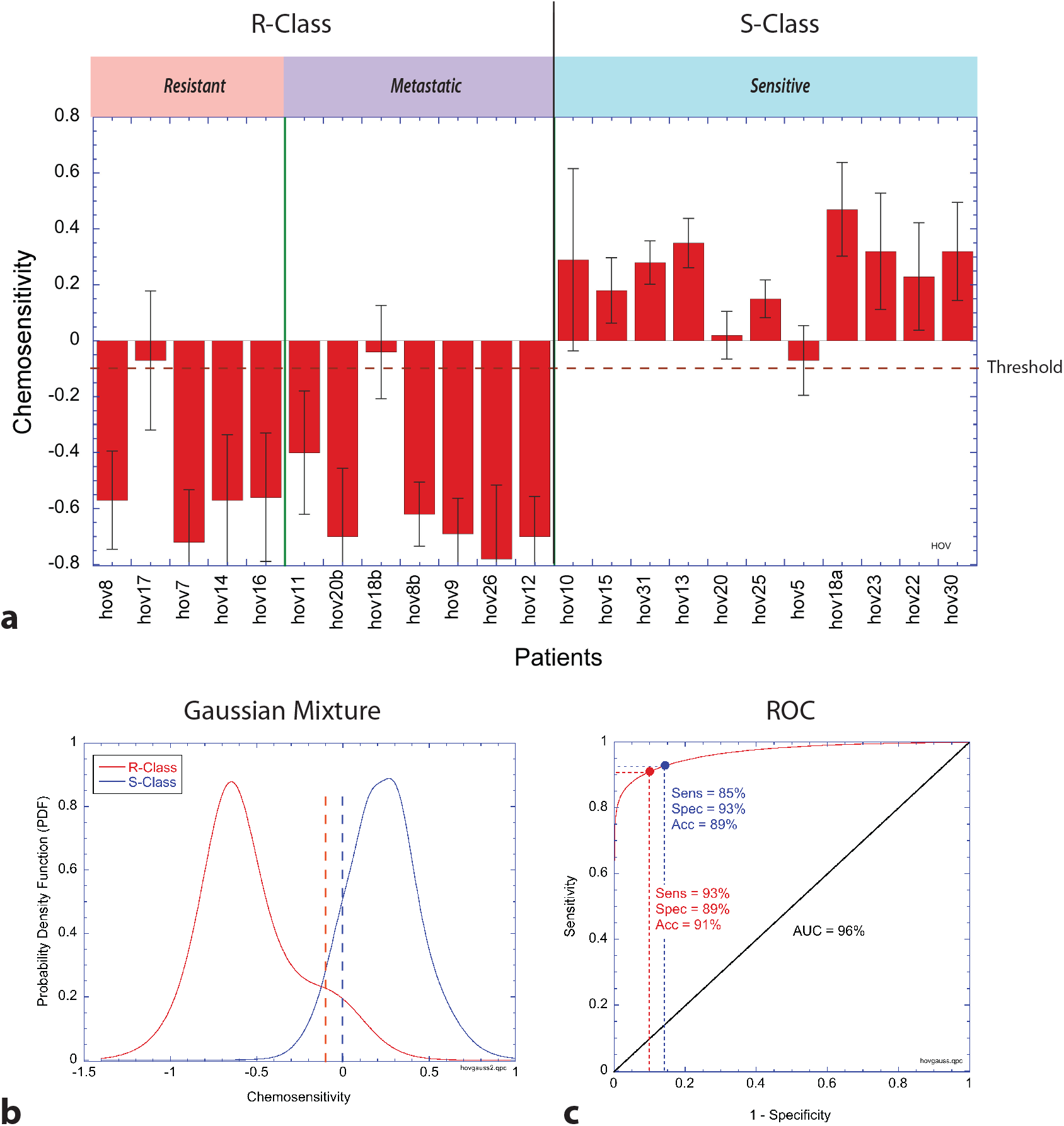
Chemosensitivity prediction with hold-out cross-validation for the training-set samples using an ensemble of algorithms trained on samples immobilized using poly-lysine. a) Ensemble predicted chemosensitivity. Error bars are standard errors on the ensemble averages (calculated from Fig. S4). R-Class are the resistant and metastatic specimens, and S-Class are the sensitive specimens. b) Gaussian mixture model of the chemosensitivity prediction probability density functions (PDF) for R-Class and S-Class specimens. The vertical dashed lines represent two possible decision points: a pre-fixed threshold at zero (blue) and for optimum sensitivity and specificity performance (red). c) Receiver operator curve (ROC) by integrating the Gaussian PDFs. The two decision points are shown. The prediction accuracy is approximately 90% when distinguishing between the two phenotypic signatures.

A continuous-valued probability distribution function (PDF) is generated from the means and standard errors by generating a Gaussian mixture model composed of Gaussian probabilities assigned the mean and standard deviation from each patient. The resulting PDF is shown in **Fig. 3b** showing a clear separation between the R-class and the S-class patients. The mean separation is 0.77 with confidence interval (0.59 – 0.95). The receiver operating characteristic (ROC) curve is generated from the Gaussian mixture model and is shown in **Fig. 3c** with an area under the curve (AUC) of 97%. By using the continuous-valued Gaussian mixture to generate the ROC instead of the discrete values, the ROC is less sensitive to threshold selection. Two decision points (thresholds) on the ROC are shown that relate to two thesholds: one (blue) at the preset threshold between positive and negative prediction values, and the other (red) at the optimal separation between the two classes. The accuracy is approximately 90% for either decision point. For the optimal threshold, the sensitivity is 93%, the specificity is 89%, the positive likelihood ratio (PLR) is 8.6, the negative likelihood ratio (NLR) is 0.08, the positive predictive value (PPV) is 89%, and the negative predictive value (NPV) is 93%.

## Discussion

The work presented in this paper is the first application of biodynamic imaging to human tissue samples. This study included 23 specimens collected prospectively from 20 patients with ovarian cancer. Seven specimens were derived from metastatic tumors and sixteen were from primary tumors. There were three patients from which matched metastatic implant and primary tumor were collected and analyzed. All patients in the study received carboplatin-based therapy, but additional therapies administered to some patients were not tested using BDI, which is one limitation of the current study.

As shown in **Fig. 1c** and **1d** for the poly-lysine-immobilized specimens, resistant biopsies responding to the carboplatin and paclitaxel monotherapies display a spectral red shift feature that is absent from the sensitive phenotype. However, when the treatments are applied as a combination, there is broadband inhibition for both the sensitive and the resistant cohort (loss of spectral density across all frequencies represented by a blue color on the spectrogram) indicating the non-additive character of the combination therapy.

Two strong biodynamic phenotypes emerged from the analysis, as shown in the similarity matrix of **Fig. 2b** and the network of **Fig. 2c**. The resistant and the metastatic specimens share a common phenotype that is distinct from the sensitive specimens. In **Fig. 2c**, marginal members of the subgroups are the patients hov5, hov17, hov18b and hov20. These four patients share some features of both phenotypes either because they display a rare phenotype, or because of measurement error.

In **Fig. 3a**, all five ovarian tissues that had resistant clinical outcomes have resistant biodynamic signatures. Of the 7 metastatic samples, all have resistant biodynamic signatures. In the 3 cases where primary and metastatic materials were obtained from the same patient (hov8/hov8b, hov18a/hov18b and hov20a/hov20b), the metastatic tissue always displayed a resistant phenotype even when the primary tumor displayed a sensitive phenotype (hov18a/hov18b and hov20a/hov20b), although hov18b did not achieve a significant prediction. In **Fig. 3b** and **3c** the Gaussian mixture and the corresponding ROC yield an accuracy of 90% for prediction of the two phenotypes whether the threshold is fixed prior to prediction (threshold at zero) or is adjusted to optimize sensitivity and specificity. Based on these findings, a metastatic sample does not predict patient clinical outcome. Therefore, one limitation of this technique, if it is to be used to predict patient outcomes, is the necessity to acquire primary tumor tissue for the assay. When considering only the 16 primary tumor tissues in this study by excluding the metastatic specimens, the accuracy is 92% with sensitivity= 0.93, specificity = 0.88, PLR = 8.1, NLR = 0.08, PPV = 0.95 and NPV = 0.85.

The drug-response spectrograms, capturing changes in intracellular motions caused by the applied therapies, are generally able to discriminate between the two phenotypes associated with patients who were resistant or sensitive to platinum-based chemotherapy. Our findings support that BDI has the potential to predict chemotherapy outcome and warrants future testing. In ovarian cancer patients, new predictive biomarkers, such as BDI profiles, could be particularly helpful for selecting second and later lines of treatment. Because the six metastatic specimens displayed a resistant phenotype, even though the patients themselves were clinically sensitive to platinum, it is possible that metastatic implants have resistant behavior, but this possibility must be studied with a larger trial size.

## Methods

### Patient Enrollment and Sample Collection

Eligible patients were those with suspected ovarian cancer who were undergoing standard-of-care cytoreductive surgery and who were willing to allow tissues to be collected for research, if available. Forty-eight patients enrolled in the study between June 2016 and November 2018. Twenty-eight patients were withdrawn, and twenty patients who passed all selection criteria were included in the final analysis. The most common reason for withdrawal was the inability to collect sufficient tumor tissue for research at the time of surgery. A total of twenty-three biospecimens were collected and used for analysis. Of these, sixteen were primary tumors and seven were metastatic tumors. Three of the metastatic implants were collected from patients who also had primary tumors collected, allowing a direct comparison of the response of primary versus metastatic lesions to chemotherapy treatment in the chemosensitivity assay.

Patients eligible for the study were age ≥ 18 years, planning to undergo surgery or biopsy as a standard-of-care treatment for suspected ovarian cancer, with subsequent histologic confirmation of ovarian, fallopian or primary peritoneal cancer. All histological types and stages were eligible for enrollment. The study was approved by the Northwestern University Institutional Review Board (protocol # STU00202733), and all patients provided written informed consent. Tissue was deidentified before processing. Enrolled patients underwent cytoreductive surgery followed by a platinum-based chemotherapy regimen, as indicated by the treating physician, per standard of care. Patients were followed for up to 18 months for clinical outcomes. Given that most patients underwent surgery with removal of tumor bulk, response to treatment (i.e., platinum sensitivity versus resistance) was determined based on time to progression (i.e., calculated from the platinum-free interval), using standard criteria ^29^. Platinum-sensitive tumors were defined as those tumors that did not recur for ≥ 6 months, while platinum-resistant tumors were those that progressed within < 6 months after completion of platinum-based therapy. All methods were carried out in accordance with relevant guidelines and regulations. A table of enrolled patients is given in Table 1. Some patients received neoadjuvant chemotherapy prior to surgery.

**Table 1.** Enrolled patients. C = carboplatin, CT = carboplatin+paclitaxel, CGT = carboplatin+paclitaxel, gemcitabine, CG = carboplatin+gemcitabine, CD = carboplatin+docetaxel, Cyclophos = cyclophosphamide. *Patient hov7 is clear cell carcinoma which is resistant to platinum therapy.

| No. | Tissue | Pathology | Treatment | Neoadj. | Response | Cell | Immobilization |
| --- | --- | --- | --- | --- | --- | --- | --- |
| 1 | primary | papillary serous carcinoma | CT | No | sensitive | hov5 | Agar |
| 2 | primary | clear cell carcinoma | CT | No | sensitive* | hov7 |  |
| 3 | primary | serous carcinoma | CGT | Yes | resistant | hov8 |  |
| 4 | metastatic |  | CGT | Yes |  | hov8b |  |
| 5 | metastatic | serous carcinoma | CT | Yes | sensitive | hov9 |  |
| 6 | primary | serous carcinoma | CT | Yes | sensitive | hov10 |  |
| 7 | peritoneum | serous carcinoma | CT, Taxotere<br>Bevacizumab | No | sensitive | hov11 |  |
| 8 | metastatic | serous adenocarcinoma | CT, Niraparib,<br>Cisplatin | Yes | sensitive | hov12 |  |
| 9 | primary | serous carcinoma | CGT | Yes | sensitive | hov13 | poly-lysine |
| 10 | primary | clear cell adenocarcinoma | CT | No | resistant | hov14 |  |
| 11 | primary | serous adenocarcinoma | CG | Yes | sensitive | hov15 |  |
| 12 | primary | carcinosarcoma | C,<br>Pembrolizumab | Yes | resistant | hov16 |  |
| 13 | primary | serous carcinoma | CG | Yes | resistant | hov17 |  |
| 14 | primary | serous carcinoma | CG | No | sensitive | hov18a |  |
| 15 | metastatic |  | CG | No |  | hov18b |  |
| 16 | primary | serous carcinoma | CT | No | sensitive | hov20a |  |
| 17 | metastatic |  | CT | No |  | hov20b |  |
| 18 | primary | serous carcinoma | C | Yes | sensitive | hov22 |  |
| 19 | primary | serous carcinoma | CD | Yes | sensitive | hov23 |  |
| 20 | primary | endometrioid<br>adenocarcinoma | C, Cyclophos | No | sensitive | hov25 |  |
| 21 | metastatic | serous adenocarcinoma | C, Taxol,<br>Cisplatin | No | sensitive | hov26 |  |
| 22 | primary | serous carcinoma | C, Taxol | Yes | sensitive | hov30 |  |
| 23 | primary | serous carcinoma | CT | Yes | sensitive | hov31 |  |

### Sample Preparation and Immobilization

The samples were dissected to approximately 1mm^3^ pieces and immobilized in 36 wells of two 96-well plates for a total of 72 wells. Two different immobilization methods were used to keep the samples fixed during measurements. Eight samples (hov5, hov7, hov8, hov8b, hov9, and hov10, hov11 and hov12) were placed in a layer of agarose covered with culture medium, while the other samples were immobilized on poly-lysine coated plates. The switch from agar to poly-lysine was undertaken partway through this trial because poly-lysine is more effective at attaching a sample to the bottom of the plate with minimal effect on the drug response (see Fig. S1 for systematic effects of each immobilization approach). Each sample received one of four treatments: carboplatin, paclitaxel, carboplatin+paclitaxel and negative control (0.1% dimethyl sulfoxide (DMSO) in RPMI-1640 growth medium).

### Biodynamic Spectroscopy

Living biopsy materials from human epithelial ovarian cancer patients were shipped in cold-packs overnight to the measurement facilities at Animated Dynamics, Inc., where the samples were dissected into approximately 72 samples of approximately 1 mm^3^ volume and immobilized in wells of a 96-well plate. Two immobilization methods were used: soft agar (low-gel temperature agarose at 1% concentration in serum-free RPMI-1640 medium.) and poly-D lysine. Treatments of 25 µM carboplatin, 5 µM paclitaxel or 25 µM carboplatin + 5 µM paclitaxel combination were applied to individual samples in individual wells and were monitored using the biodynamic imaging system for up to 12 hours. The drug concentrations were chosen near the IC50 (in vitro 50% response) for each single-agent therapies, and these concentrations were maintained in the combination. The well replicate numbers were 17 for negative control (0.1% DMSO dimethyl sulfoxide in RPMI-1640 medium), 18 for paclitaxel, 18 for carboplatin, and 18 for carboplatin + paclitaxel. The time-course of the experiment is a 5.5-hour baseline with the system successively sampling each of 36 wells in a repeating cycle that takes 82 minutes to measure all 36 wells before repeating. The drug is administered by withdrawing approximately100 μL of old growth medium and gently pipetting 100 μL of treatment volume at twice the target concentration. The drugs are dissolved in DMSO and diluted into RPMI-1640 growth medium. The system acquires data for 12.3 hours after the drugs are applied.

### BDI Measurement and Drug Treatment

Sample imaging was carried out on the biodynamic platform (BDP) (Animated Dynamics Inc, Indianapolis). The imaging system is placed on a motorized optical platform that moves on the horizontal plane, while the plate is on a fixed mount keeping it stationary during the entire measurement. The BDI system is Mach-Zehnder interferometer using off-axis digital holography, as shown in **Fig. 1a**. The light source is a low-coherence superluminescent diode (20 mW power 50 nm bandwidth centered on 840 nm) that illuminates the sample at an oblique angle. The scattered light is collected through a Fourier imaging system that projects the Fourier transform of the tissue speckle onto the camera plane. A delay stage is placed in the reference arm to modify the optical path length of the arm to achieve depth-selective coherence gating of the sample. The optical tissue section is reconstructed through a 2D FFT of the digital hologram. The frame rate of the camera is 25 frames per second with a Nyquist sampling frequency of 12.5 Hz. The time series of the intensity fluctuations are Fourier transformed to a frequency spectrum which is averaged over the sample. When a drug is applied to the tissue, the motions change, which are captured by shifts in the Doppler spectrum. These shifts are represented as drug-response spectrograms that track shifts in spectral density as a function of time.

The drug-response spectrograms are generated by creating a series of logarithmic fluctuation power spectra at successive times and subtracting the average baseline. The resulting spectrograms display the shift in Doppler spectral content of the sample over the 17.8-hour assay. Details of tissue dynamic spectroscopy are given in previous publications^19, 24^. The baseline spectrum *S*(ω,0) is defined as the last 4 loops prior to the treatment. The drug-response spectrogram is defined as *D*(*ω, t*) = *log S* (*ω, t*) − *log S* (*ω*, 0), where the time index *t* represents the loop number. A spectrogram is generated for each well for a given patient and a given treatment. An algorithm assigns a data quality factor for each well based on multiple quality control criteria such as a sudden jump in brightness/intensity (indicating immobilization shift) or low cell activity. The value is initialized at unity and is reduced by a factor of 2 for each violation of a criterion. Typical data qualities are around 0.25. The spectrograms are averaged over the replicate numbers for the treatment weighted by the DQ. A patient is thus represented by an average spectrogram for each treatment.

### Biomarker Definition and Feature Extraction

There are approximately 5 spectral bands that can be defined in the drug-response spectrograms. These are defined in **Table 2** with their presumed biophysical origins (See Ref. 9 for biophysical origins). The time-frequency representation of the drug-response spectrograms are converted into 18 quantitative biomarkers through linear filters. Additional biomarkers represent sample preconditions and changes in these conditions caused by the applied drug.

**Table 2:** Spectral Bands (Backscattering geometry with λ = 840 nm)

| Band Name | Frequency Range | Speed Range | Biophysics Origins |
| --- | --- | --- | --- |
| Cell Motility Band | 10 mHz | < 3 nm/s | Crawling |
| Rheology Band | 12.5 mHz – 100 mHz | 4 nm/s – 30 nm/s | Shape change |
| Mid Band | 100 mHz – 1 Hz | 30 nm/s – 300 nm/s | Membrane/Nuclear |
| High Band | 1 Hz – 10 Hz | 300 nm/s – 3 μm/s | Organelle transport |
| Nyquist Band | 12.5 Hz | > 4 μm/s | Vesicle transport |

The time-frequency spectrograms are converted into feature vectors with parts or patterns of the spectrograms. In addition to spectrogram-based features, there are also preconditions (such as sample brightness and dynamic range, etc.) as well as drug-induced changes in these preconditions. All the raw biomarkers are defined in **Table S3**. The time-frequency decomposition is approached globally and locally. Global patterns are generated as low-order Legendre polynomials. These polynomials are taken as an inner product over the spectrograms to generate Legendre coefficients that represent the global features of the spectrograms. Orders 0, 1 and 2 are used along the frequency and time axes to generate 9 global features. Local patterns are simply low, mid, and high-frequency bands with average, linear and quadratic time dependence for 9 local features. The precondition biomarkers are NSD, BSB, NCNT, DR, NY, KNEE, HW, S, SF that represent properties such as sample speckle contrast, brightness, sample size, dynamic range of the signal, the spectral density of the Nyquist floor, the knee frequency, the half-width of the Doppler spectrum, and the slope of the roll-off above the Doppler knee frequency, respectively. (Note that NCNT, KNEE, and S are subject to fitting errors and are down-selected as features.) Each precondition is changed by the drug treatment, providing additional features that are the changes in the preconditions from baseline to endpoint of the assay. There are 27 drug-response features for a given drug: 18 are based on spectrograms and 9 are drug-induced changes in preconditions. These 27 features are concatenated for each drug to create a feature vector of 27×3 = 81 drug elements. The 9 global and 9 local time-frequency filter masks are shown in **Fig. 4a)** and **4b**, respectively. The decomposition into the 18 spectrogram-based biomarkers generates strong covariance among the biomarkers. Therefore, singular-value decomposition (SVD) is used to find orthogonal time-frequency biomarkers that are linear combinations of the original biomarkers. Linear combinations of the spectrogram-based biomarkers with the largest signal-to-noise (also called the z-factor, which is the mean difference divided by the standard deviation) for the resistant/sensitive groups are shown in **Fig. 4c**. The biomarkers are designated by the level of the orthogonal component and by the treatment. For instance, BM7tax+carb represents the 7th principal component in the singular-value decomposition of the drug response under Taxol+carboplatin treatment.

**Figure 4.**
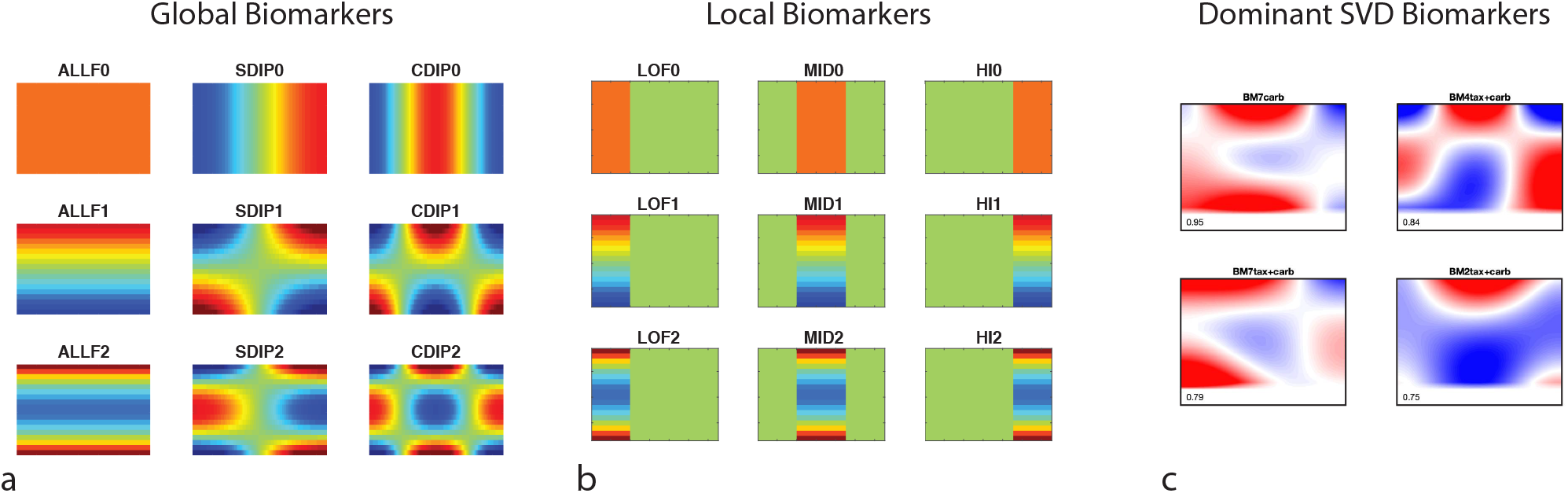
Time-frequency biomarker masks. a) Global biomarker filters are low-order Legendre polynomials along the time and frequency axes. b) Local biomarker filters are low, mid- and high-frequency bands with 0, 1 and 2^nd^-order polynomial time dependence. c) The 4 dominant singular-value decomposition (SVD) biomarkers represented as time-frequency patterns. The numerical values are the z-factors for each biomarker.

### Machine Learning Methods

The similarity matrix measure that we selected is correlation contrast that equals the correlation coefficient when the magnitudes of the vectors are similar, but down-weights the correlation if there is a mismatch in vector magnitude. This down-weights similarities when the amplitudes of the two feature vectors are not matched. The motivation for this metric is the fringe contrast in interferometric measurements. The metric is

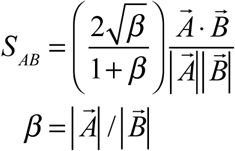

where *β* is the amplitude ratio of the two vectors. This metric is used to construct the similarity matrix in **Fig. 2b**. The similarity matrix is the central data structure for all down-stream machine learning methods. For instance, the similarity network in **Fig. 2c** is constructed using a k-neighbor algorithm that connects each node to k nodes to which it shares the highest similarity. The network in **Fig. 2c** used k = 3.

The ensemble approach to patient chemosensitivity prediction used 4 binary classifiers and averaged the results: 1) a single-neuron perceptron (logistic regression), 2) a continuous-valued recurrent neural network, 3) log-likelihood and 4) a binary network analysis. Logistic regression minimizes the cost function ^30^ using ridge regularization

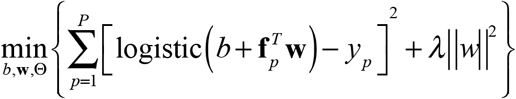

where y_p_ = ±1 is the objective classification of the p-th patient, **f**^T^_p_ is the feature vector for the p-th patient, b and **w** are adjusted to minimize the squared error, and λ = 0.1 is the regularization parameter. Once the parameters are trained, the predicted chemosensitivity P_j_ of hold-out or non-training feature vectors are predicted as

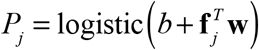

The continuous-valued recurrent neural network uses Gibbs sampling ^31^ and sequential updating of the j-th feature vector from the set of inner products where the probability of selecting the p-th feature vector as the update is

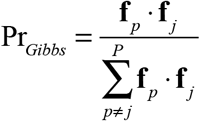

and where the updating is performed at each iteration on a single randomly-selected feature of the p-th feature vector. The sampling is iterated to convergence to the updated feature vector **f**’_j_, and the similarity of the updated feature vector is calculated to each of the training features and used as the weighting factor on the objective classification y_p_. The predicted chemosensitivity P_j_ of hold-out or non-training feature vectors are predicted as

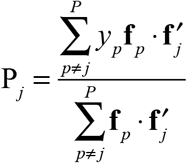

The log-likelihood prediction constructs a Gaussian mixture model to construct probability distribution functions L_-1,1_ of the features among the training set ^31^. In the binary classifier, each PDF has a classification index. The predicted chemosensitivity P_j_ of hold-out or non-training feature vectors are predicted as

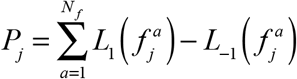

The network binary classifier uses the adjacency matrix A(j,l) constructed by the k-neighbor algorithm from the similarity matrix. The predicted chemosensitivity P_j_ of hold-out or non-training feature vectors are predicted as

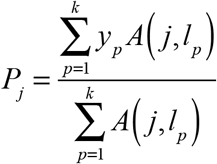

where l_p_ is the target node of the p-th link to the j-th node. The ensemble learning takes the ensemble average over the four binary classifiers to yield the average predicted chemosensitivity of **Fig. 3**. The error bars are the standard error of the ensemble values added in quadrature for the root-variance of the training sub-sets of **Fig. S4**.

## Acknowledgements

This work was supported by grants NSF 1911357-CBET and NIH NIBIB 1RO1EB016582 and by the Purdue Cancer Center.

## Author contributions statement

Z.L. and R.A. contributed equally to the data collection. Z. L. and D. N performed the data analysis and wrote the main manuscript text and prepared the figures. J.T. provided the sample handling protocols for biodynamic imaging. W. S., M. K., N.N. and D.M. provided all clinical samples, performed clinical data collection and analysis. All authors reviewed and edited the manuscript.

## Data Availability

The data and codes that support the findings of this study are available <u>at GitHub from</u>https://github.rcac.purdue.edu/nolte/SciRep2020-Data-and-Files or from the corresponding author upon reasonable request.

