## Supplemental Information for "Intracellular Optical Doppler Phenotypes of Chemosensitivity in Human Epithelial Ovarian Cancer"

### Patient Characteristics

The patient characteristics and statistics for the trial enrollments are shown in **Table S1**.

**Table S1.** Patient characteristics

| **Patient Characteristics** | **Frequency (n=20) [%]** |
|---|---|
| Disease Status | |
| Newly Diagnosed | 18 [90%] |
| Platinum-Sensitive Recurrent Disease | 2 [10%] |
| **Diagnosis** | |
| Ovarian Cancer | 17 [85%] |
| Fallopian Tube Cancer | 2 [10%] |
| Primary Peritoneal Cancer | 1 [5%] |
| **FIGO Tumor Stage** | |
| I/II | 5 [25%] |
| III/IV | 15 [75%] |
| **Tumor Grade** | |
| High Grade or Poorly Differentiated | 19 [95%] |
| Low Grade or Well Differentiated | 1 [5%] |
| **Tumor Histology** | |
| Serous | 16 [80%] |
| Clear Cell | 2 [10%] |
| Endometrioid | 1 [5%] |
| Carcinosarcoma | 1 [5%] |
| **Cytoreduction** | |
| Optimal Primary Cytoreduction | 16 [80%] |
| Suboptimal Primary Cytoreduction | 2 [10%] |
| Secondary Cytoreduction | 2 [10%] |
| **Systemic Therapy prior to Specimen Collection** | |
| None | 8 [40%] |
| Neoadjuvant Platinum + Taxane | 10 [50%] |
| Multiple Prior Therapies, including >=1 Platinum | 2 [10%] |
| **System Therapy after Specimen Collection** | |
| Platinum + Taxane | 12 [60%] |
| Platinum + Gemcitabine | 5 [25%] |
| Platinum + Cyclophosphamide | 1 [5%] |
| Platinum + Immunotherapy | 1 [5%] |
| Platinum Monotherapy | 1 [5%] |
| **Platinum Free Interval (PFI) (Duration of Remission after Platinum Therapy)** | |
| PFI ≥ 6 months (Platinum-Sensitive) | 16 [80%] |
| PFI < 6 months (Platinum-Resistant) | 4 [20%] |
| **Other Patient Demographics** | **Median [Range]** |
| Age | 63.5 [47-76] |
| Number of Prior Systemic Therapies | 1 [0-5] |
| Number of Prior Platinum Therapies | 1 [0-3] |

### Immobilization

The two different sample immobilization methods were used. The first 8 samples were immobilized using agar, and the remaining 15 samples were immobilized using poly-lysine. The shift occurred because poly-lysine was found to provide better sample stabilization. However, this shift created systematic differences in some BDI features. There is a difference in the means for values of baseline biomarkers like NSD in a two-sample t-test for samples immobilized with agarose vs with poly-lysine. The difference in drug response is significant for paclitaxel and its combination with carboplatin, but for not carboplatin only. D'Agostino-Pearson normality tests were used to validate the t-test normalized data assumption. (p-values are given in Table S2). The low NSD values found in poly-lysine immobilized samples indicate that poly-lysine is more effective for sample attachment. Drugs containing paclitaxel have lower SDIP0 values that may indicate that the paclitaxel mechanism of action, i.e. targeting tubulin and stabilizing the microtubule polymer, may be interacting with the mechanical properties of the agarose, creating trends that show up as part of the drug responses. The comparison of agar to poly-lysine is shown in Fig. S1. Because of this immobilization systematic, the primary analysis trains exclusively on the 15 samples immobilized by poly-lysine, then uses the trained model to test the agar samples and the metastatic samples (of either immobilization). This approach down-weights the feature selection that might be influenced by the agar mechanical properties.

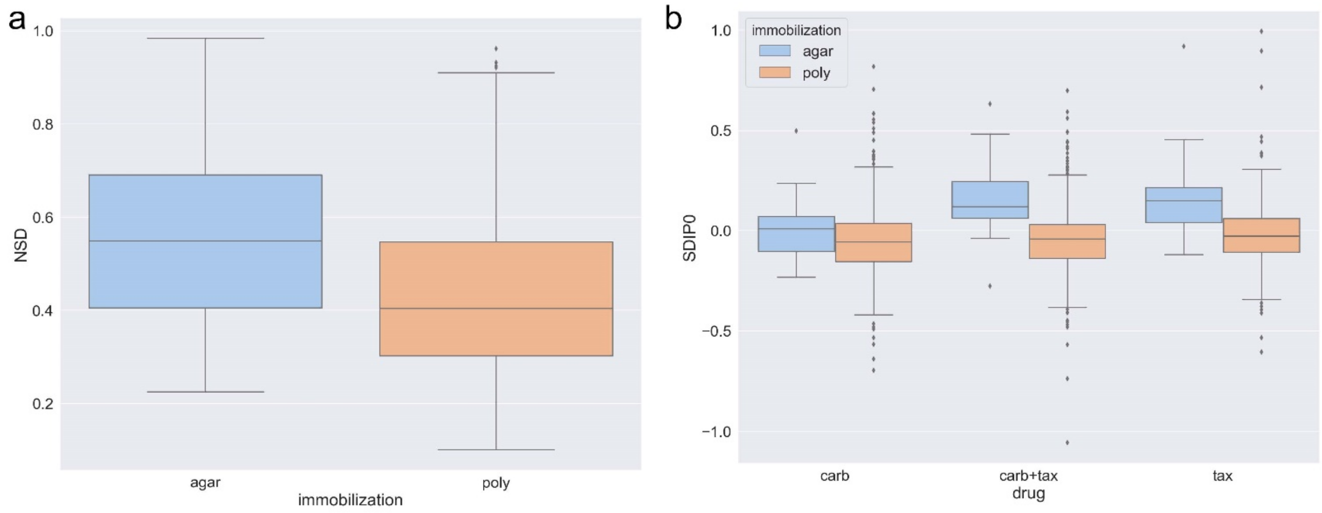

**Figure S1:** comparison of feature value from samples under different immobilization methods. a) NSD values. b) SDIP0 values for three treatments

**Table S2**: p-values for statistical tests used to study agar versus poly immobilization

| | NSD | SDIP0 (carb) | SDIP0 (carb+tax) | SDIP0 (tax) |
|---|---|---|---|---|
| t-test for agar/poly-lysine comparison | $2\times10^{-15}$ | 0.06 | $7\times10^{-11}$ | $2\times10^{-7}$ |
| Normality test for agar samples | $5\times10^{-5}$ | 0.008 | 0.04 | $2\times10^{-8}$ |
| Normality test for poly-lysine samples | $7\times10^{-10}$ | $2\times10^{-7}$ | $10^{-5}$ | $4\times10^{-20}$ |

### Biomarker Definition

Table S3 is a comprehensive description of all 40 metrics, or features, associated with a patient and drug. The first 9 are the global biomarkers, and the next 9 are the local biomarkers, discussed in the main text. These 18 are all based on the time-frequency format of the drug-response spectrogram. The frequency bands are described in Table 2 in the main text. The time dependence is simple polynomial: 0 is constant, 1 is linear, and 2 is quadratic.

The biomarkers 19 – 27 are drug-induced changes in the preconditions 28 – 36. NSD is the normalized standard deviation, also known as temporal speckle contrast. BSB is the brightness of the sample. NCNT is the number of pixels in the cross-sectional image of a target. DR is the “vertical” dynamic range of the spectral density of a power spectrum. NY is the value of the spectral density at the Nyquist frequency. KNEE is the knee frequency of the fluctuation spectrum at which the power falls to half of its low-frequency value. HW is the half-width of the spectrum, closely related to KNEE. S is the spectral slope (linear on log-log) of the power spectrum for frequencies above the knee frequency and is closely related to SF which uses a nonlinear fitting method to measure the slope. The final metrics in Fig. S3 include three measures of the baseline B0, B1 and B2 which each represent constant, linear and quadratic frequency dependence. The final metric DQ is the data quality assigned to each well or to each patient and drug.

**Table S3: Definitions of Biodynamic Biomarkers**

| | Biomarker Name | Description |
|---|---|---|
| | | **Global Spectral Biomarkers** |
| 1 | ALLF0 | All frequencies. All times |
| 2 | SDIP0 | Blue shift: All times |
| 3 | CDIP0 | Middle-out: All times |
| 4 | ALLF1 | All frequencies. Linear time dependence |
| 5 | SDIP1 | Blue shift: Linear time dependence |
| 6 | CDIP1 | Middle-out: Linear time dependence |
| 7 | ALLF2 | All frequencies. Quadratic time dependence |
| 8 | SDIP2 | Blue shift: Quadratic time dependence |
| 9 | CDIP2 | Middle-out: Quadratic time dependence |
| | | **Local Spectral Biomarkers** |
| 10 | LOF0 | Low-frequencies: All times |
| 11 | MID0 | Mid-frequencies: All times |
| 12 | HI0 | Hi-frequencies: All times |
| 13 | LOF1 | Low-frequencies: Linear time dependence |
| 14 | MID1 | Mid-frequencies: Linear time dependence |
| 15 | HI1 | Hi-frequencies: Linear time dependence |
| 16 | LOF2 | Low-frequencies: Quadratic time dependence |
| 17 | MID2 | Mid-frequencies: Quadratic time dependence |
| 18 | HI2 | Hi-frequencies: Quadratic time dependence |
| | | **Change in Precondition** |
| 19 | DNSD | Change in normalized standard deviation (NSD) |
| 20 | DBSB | Change in back-scatter brightness (BSB) |
| 21 | DNCNT | Change in number of pixels (NCNT) |
| 22 | DDR | Change in dynamic range (DR) |
| 23 | DNY | Change in Nyquist floor (NY) |
| 24 | DKNEE | Change in knee frequency (KNEE) |
| 25 | DHW | Change in half-width (HW) |
| 26 | DS | Change in Slope (S) |
| 27 | DSF | Change in linear slope (SF) |
| | | **Precondition** |
| 28 | NSD | Normalized standard deviation (NSD) |
| 29 | BSB | Back-scatter brightness (BSB) |
| 30 | NCNT | Number of pixels (NCNT) |
| 31 | DR | Dynamic range (DR) |
| 32 | NY | Nyquist floor (NY) |
| 33 | KNEE | Knee frequency (KNEE) |
| 34 | HW | Half-width (HW) |
| 35 | S | Slope (S) |
| 36 | SF | Linear slope (SF) |
| 37 | B0 | Baseline: all frequencies |
| 38 | B1 | Baseline: linear frequency |
| 38 | B2 | Baseline: quadratic frequency |
| 40 | DQ | Data Quality |

The biomarkers in Table S3 have a covariance matrix with off-diagonal values that measure the correlations among them. Therefore, we use principal component analysis (PCA) based on singular vector decomposition (SVD) to pool the biomarkers into a smaller number of independent biomarkers. The feature selection is described in detail in the main text. The four features selected in this study are shown in Table S4 along with the coefficients for each of the raw features in Table S3.

**Table S4. Coefficients of Raw Features combined into Principal Components**

| | **BM7C** | **BM2TC** | **BM4TC** | **BM7TC** |
|---|---|---|---|---|
| ALLF0 | 0.3359 | -0.209 | 0.0488 | 0.2833 |
| SDIP0 | -0.2051 | -0.0079 | 0.1717 | -0.3371 |
| CDIP0 | 0.2623 | 0.0497 | -0.0166 | -0.1102 |
| ALLF1 | -0.0686 | 0.386 | 0.0133 | 0.0016 |
| SDIP1 | 0.0355 | -0.0034 | -0.2491 | -0.0186 |
| CDIP1 | 0.135 | 0.3571 | 0.5599 | 0.1908 |
| ALLF2 | 0.324 | 0.2502 | -0.1074 | 0.3021 |
| SDIP2 | -0.0968 | 0.069 | 0.042 | -0.3345 |
| CDIP2 | 0.308 | 0.0499 | 0.3578 | 0.0476 |
| LOF0 | 0.2654 | -0.0776 | -0.1592 | 0.4261 |
| MID0 | 0.3493 | -0.1595 | 0.0381 | 0.1936 |
| HI0 | 0.2201 | -0.2164 | -0.0111 | 0.1721 |
| LOF1 | -0.1274 | 0.1884 | -0.1198 | -0.0806 |
| MID1 | -0.0123 | 0.408 | 0.1757 | 0.0741 |
| HI1 | -0.1945 | 0.3733 | -0.5031 | -0.1617 |
| LOF2 | 0.2112 | 0.063 | -0.0942 | 0.3908 |
| MID2 | 0.3301 | 0.2206 | -0.0034 | 0.2624 |
| HI2 | 0.1719 | 0.3395 | -0.0363 | -0.031 |
| DNSD | 0.0361 | -0.0292 | 0.0128 | 0.0494 |
| DBSB | 0.0084 | 0.041 | 0.0246 | -0.0043 |
| DNCNT | -0.0001 | -0.0004 | 0 | 0.0003 |
| DDR | 0.0553 | 0.0269 | -0.0294 | 0.1008 |
| DNY | 0.0434 | -0.043 | -0.0396 | 0.0302 |
| DKNEE | 0.0008 | 0 | -0.0001 | 0.0011 |
| DHW | 0.0476 | 0.0314 | -0.0282 | 0.0256 |
| DS | 0.0004 | -0.0003 | -0.0001 | 0.0011 |
| DSF | 0.0561 | -0.0122 | -0.0124 | 0.1204 |

### Spectral Response to Refreshed Medium

The growth medium is RPMI-1640, and the carrier for adding drugs to the medium is 0.1% DMSO. Therefore, 17 replicates of the negative control (0.1% DMSO in RPMI-1640 medium) are applied for each patient to measure the response of the living tissue to the refreshed medium that contains fresh nutrients and oxygen. The spectrogram of the negative control is shown in **Fig. S2** averaged over all replicates over all patients. The baseline is established for 4 hours and the medium is applied. The biopsy samples respond to the refreshed medium with a broad-band increase in spectral density centered around 0.1 Hz (average intracellular speed 300 nm/sec). This enhanced activity is a general property of living samples responding to refreshed medium that has been observed across multiple disease types. Because this non-negligible response is part of every drug signature, this background spectrogram is subtracted from the drug responses on a patient-by-patient basis. The spectrograms shown in the figures in the main text are all after this background subtraction.

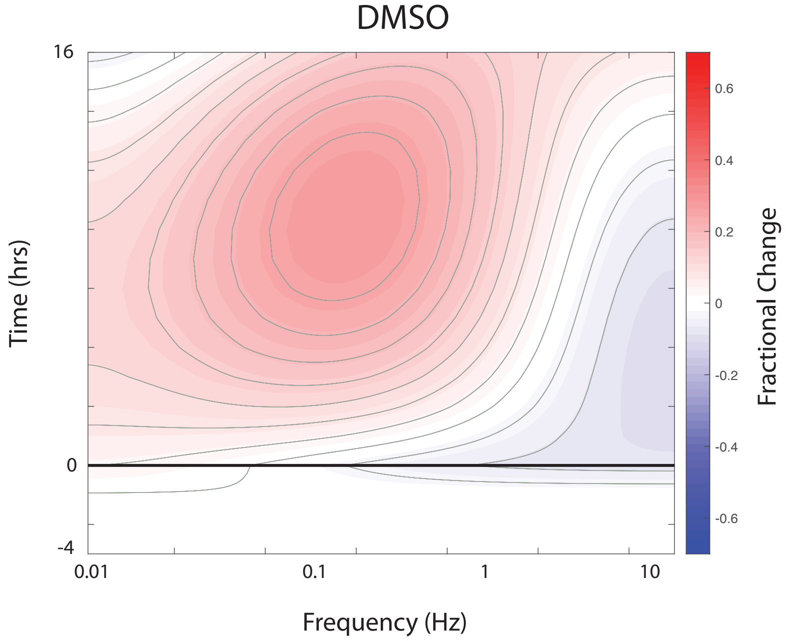

**Fig. S2** Average spectrogram response to the negative control (0.1% DMSO in RPMI-1640 growth medium). The biopsy samples respond with increased spectral weight across nearly the full frequency band. This pattern is consistent with increased cellular metabolism in response to the refreshed medium. This background response is subtracted from each drug response.

Sample-to-sample variance is a key aspect of live-tissue measurements caused by sample heterogeneity. An important distinction is well-to-well variability among spectrograms for a given patient, compared to the patient-to-patient variability. The first is analogous to homogeneous broadening, and the second is analogous to inhomogeneous broadening. Standard deviations of the spectrograms are shown in **Fig. S3**. The average patient spectrogram standard deviation is shown in **Fig. S3a**, and the average standard deviation across all patients is shown in **Fig. S3b**. The peak standard deviation in the latter case is 0.4 about 8 hours after the medium refresh and in the former is 0.27. Therefore, there is more variance patient-to-patient than well-to-well for a given patient. The standard deviations for the R-class and the S-class are given in **Figs. S3c** and **S3d**. The S-class shows larger variability among patients than the R-class. It is important to note that with 18 well-replicates per treatment, the maximum standard error on a drug-response spectrogram is approximately 0.1, or a ±10% change in spectral density at low frequencies. The standard deviation at the Nyquist floor is much smaller, which suggests that mid and high frequencies may be more reliable as biomarkers than lower frequencies. The maximum standard deviation for a given patient is 0.27 at low frequencies approximately 8 hours after treatment. With 18 replicates per treatment per patient, this represents ±7% spectral density uncertainty at that time and frequency. The average standard deviation over the entire time-frequency plane is 0.15 and the average standard error on 18 replicates is then about 4%. This 4% change in spectral content is then the detection limit of a drug effect for a given patient.

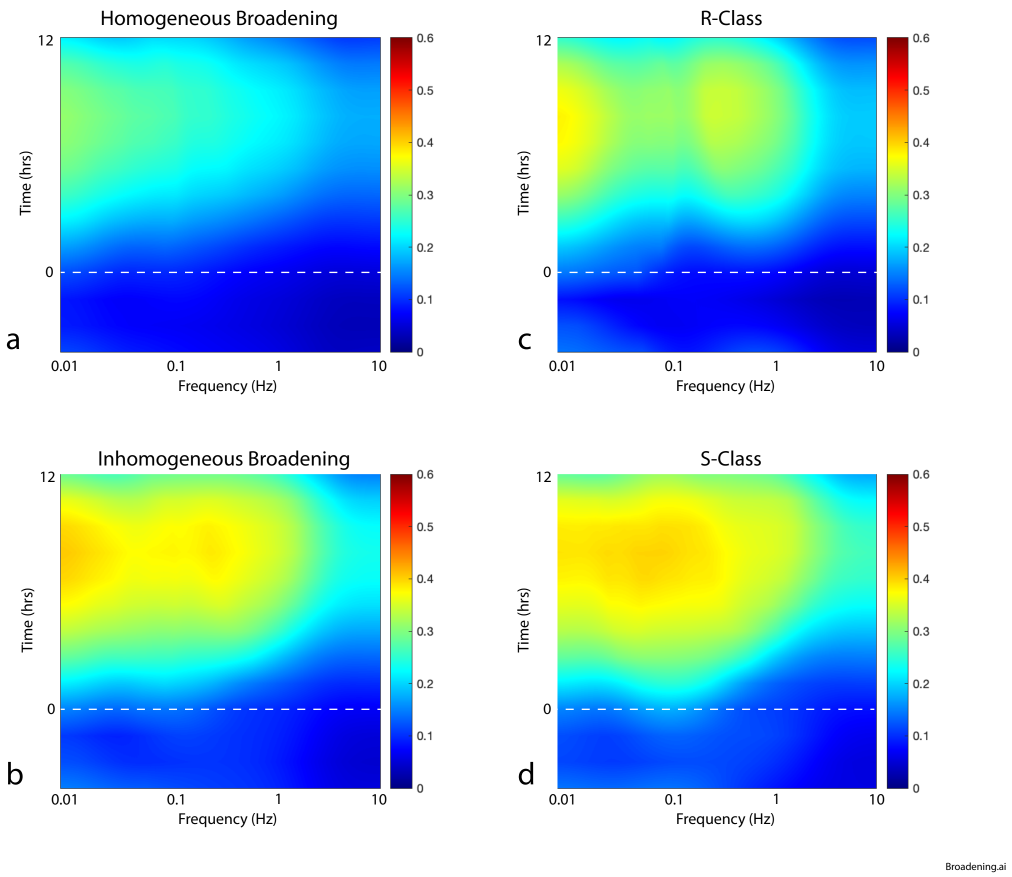

**Fig. S3.** Homogeneous versus inhomogeneous broadening of negative control spectrograms. a) Patient-based average standard-deviation of neg-control. This is the average well-to-well variability for a patient and represents the homogeneous broadening. b) Standard deviation of all negative control wells. This is the patient-to-patient variability and represents inhomogeneous broadening. c) Standard deviation of R-class negative control wells. d) Standard deviation of S-class negative control wells.

### Training-Set Stability for Predicting Chemosensitivity

The chemosensitivity values for each patient in the study, presented in **Fig. 3a** in the main text, is based on a training set for only poly-lysine immobilization of sensitive and resistant primary tumor biopsies. The metastatic samples (hov8b, 9, 11, 12, 18b, 20b, 26), as well as the agar-immobilized samples (hov5, 7, 8, 10), were then predicted using the trained algorithm. Furthermore, the poly-lysine-immobilized training set was predicted using one-hold-out.

This training subset was chosen because of the stability of the poly-lysine immobilization. However, other training subsets are possible. For instance, one could train on all the samples and predict chemosensitivity using one-hold-out for each one. The results are shown in **Fig. S4** as the red bars (error bars are the standard error on the average of the ensembles). Alternatively, the training set can be the agar and poly-immobilized samples, predicted using hold-out, and predicting the metastatic samples using the trained algorithm. The results are shown as the green bars in **Fig. S4**. These are reasonably disparate choices for the training set, and most of the patients share similar chemosensitivity values among all three training methods. Notable exceptions are patients hov17, hov11, hov18b, hov10 and hov20. Therefore, 84% of the patients predict consistently among the different training subset methods.

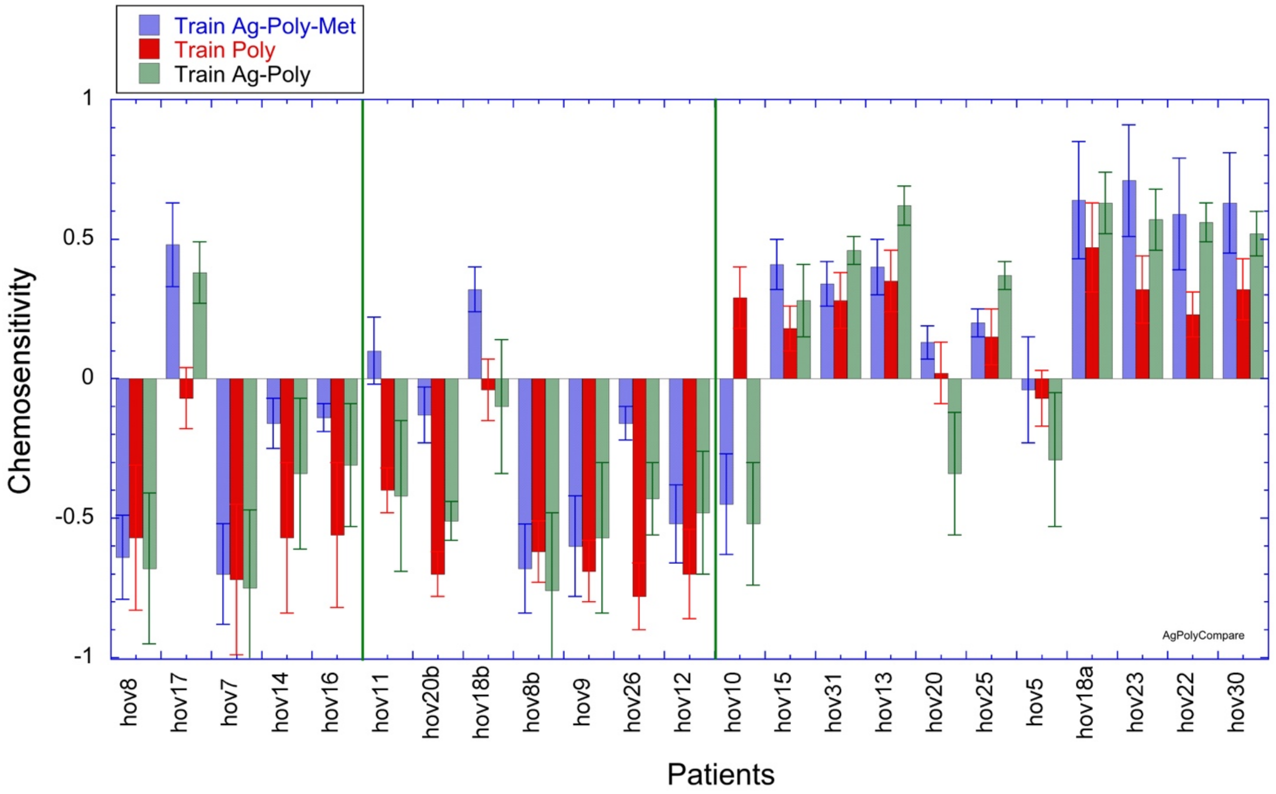

**Fig. S4** Comparison of three alternate training methods to measure patient chemosensitivity. The red bars are when training using the full set of agar-immobilized, poly-immobilized and metastatic samples and predicting using one hold out. The blue bars are when training only on the poly-immobilized samples and testing both agar and metastatic samples (the same as **Fig. 3** in the manuscript). The green bars are when training using both agar and poly immobilization, predicting the training patients using one hold out, and predicting the metastatic patients using the trained classifier. The error bars are the error on the mean from the ensemble of hold-out prediction algorithms. The patients who have inconsistent results among the different approaches are hov17, hov11, hov18b, hov10, hov20 and hov5. Four of these patients are the ones with weak predictions from **Fig. 3** in the main text.

The statistical analysis of the poly-immobilized predictions are given in **Table S5** based on the decision point that optimizes the sensitivity and specificity.

**Table S5. Statistics for Assay Performance**

| Mean R | Mean S | Std R | Std S | H | p-value |
|---|---|---|---|---|---|
| -0.5350 | 0.2309 | 0.2454 | 0.1540 | 1 | < 0.001 |

| Max CI | Min CI | Mean Diff | t-statistic | Mut. Info. | z-factor |
|---|---|---|---|---|---|
| 0.9456 | 0.5862 | 0.7659 | 8.9 | 5.2 | 1.8 |

| AUC | ACC | Sens | Spec | PPV | NPV |
|---|---|---|---|---|---|
| 0.96 | 0.91 | 0.93 | 0.89 | 0.89 | 0.93 |

### Original Spectrograms and Features

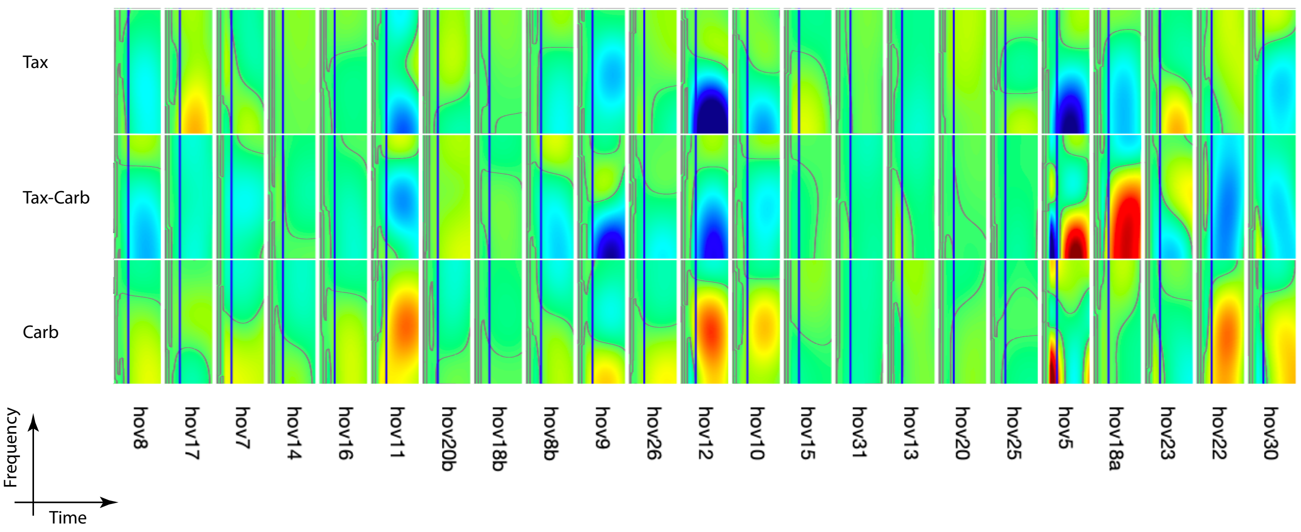

**Fig. S5** Original DMSO-subtracted spectrograms for all 23 specimens for carboplatin, Taxol and carboplatin+Taxol. The axes orientations have been switched for ease of viewing. Color scale is -60% to +60%

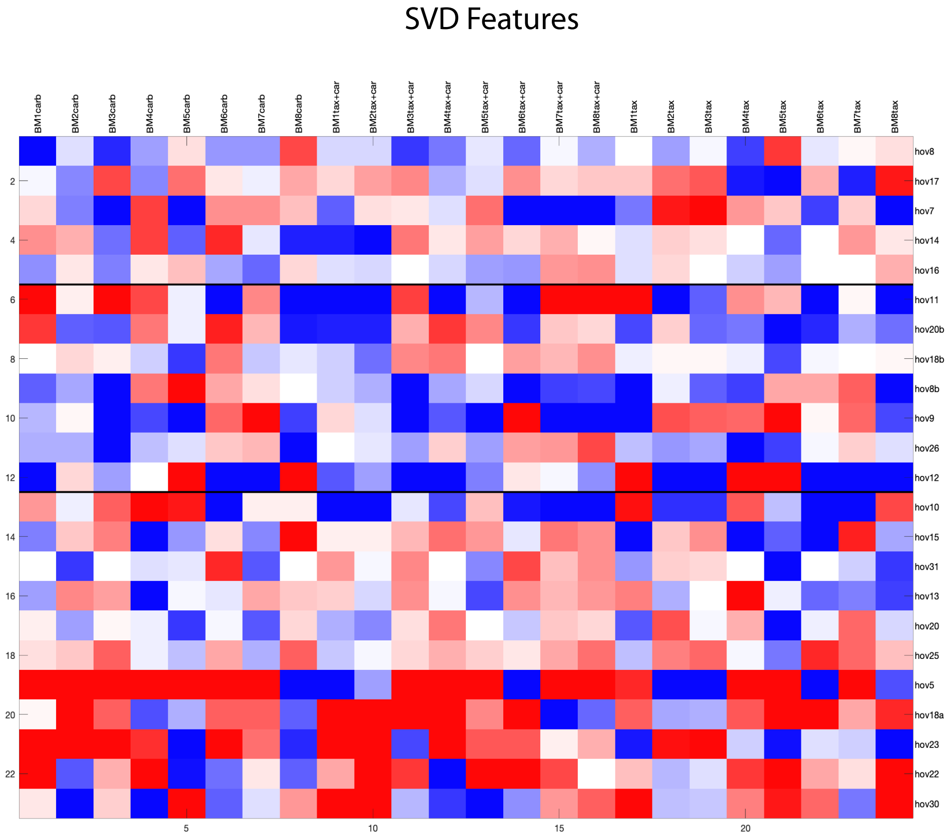

**Fig. S6** All SVD feature vectors. The features with the highest z-factors are selected for downstream analysis.

### Comparison of Human/Mouse/Cell-Line Drug Responses

Our previous work (Ref. 23) studied human ovarian cell lines grown as spheroids or as mouse xenografts. The response to carboplatin for sensitive (A2780) and resistant (CP70) cell lines are shown in **Fig. S7** compared to the results from the current work on the human biopsies. The biopsies share similarities with the spheroids, but not the mouse explants. The spheroids and biopsies have only human ovarian constituents, while the explants have constituents from the mouse host (stroma, fibroblasts and possibly immune cells), which may contribute to the differences.

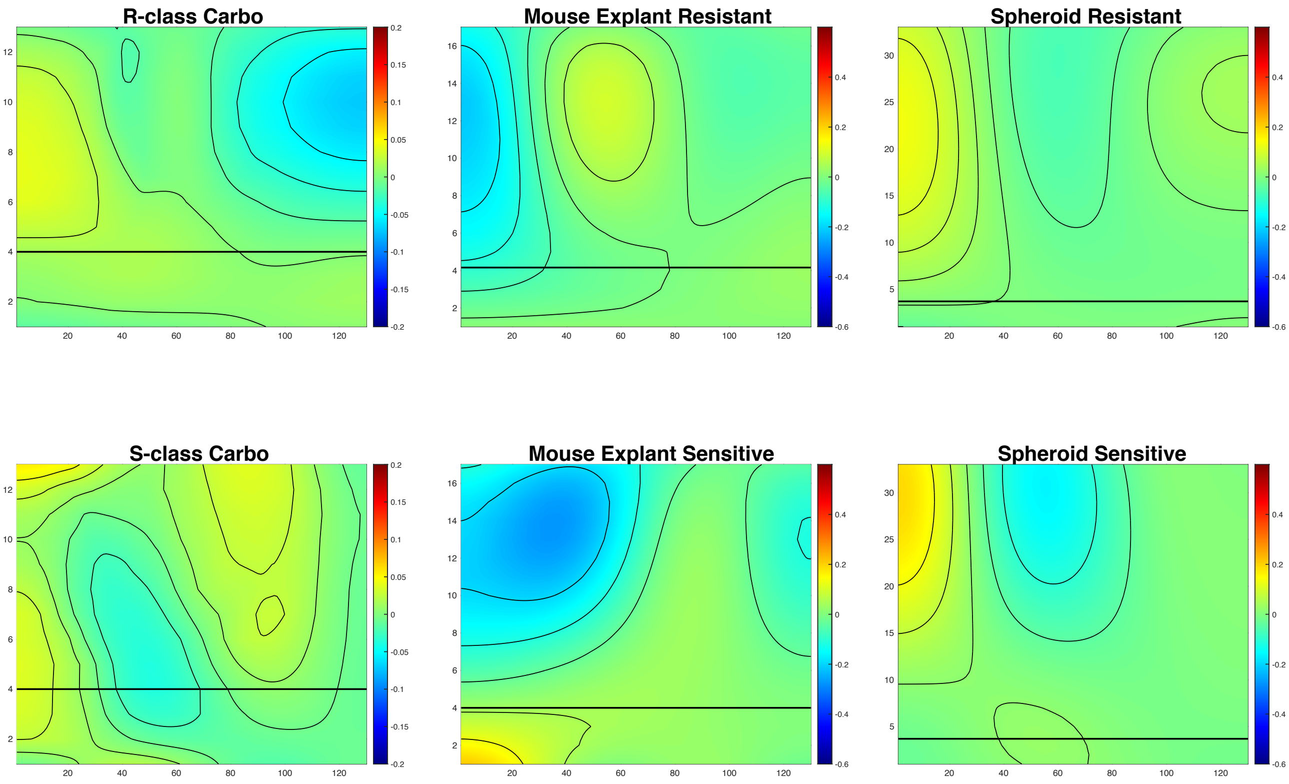

**Fig. S7** Comparison of averaged spectrograms of three ovarian tissue types for 10 μM carboplatin treatment. The explants are mouse xenografts grown from A2780 (sensitive) and CP70 (resistant) cell lines. The spheroids are grown in bioreactors from the same cell lines. The biopsies are the human biopsies from this study. The human biopsies show similar responses to the spheroids but not the mouse explants.
